# Turn-taking-like temporal coordination of ultrasonic vocalizations during close-range social interactions in Mongolian gerbils (*Meriones unguiculatus*)

**DOI:** 10.64898/2026.05.28.728373

**Authors:** Ryo Nishibori, Naoki Matsumoto, Yume Kinoshita, Kazuki T. Shin’ya, Yuki Ito, Yuta Tamai, Kohta I. Kobayasi

**Author notes:** Department of Physiology, Jichi Medical University, Tochigi, Japan (Ryo Nishibori). Department of Psychology, Keio University, Tokyo, Japan (Kazuki T. Shin’ya). Faculty of Sociology, Yamato University, Osaka, Japan (Yuki Ito). ***Corresponding authors*** Ryo Nishibori & Kohta I. Kobayasi, RN, KIK.

## Abstract

Vocal communication depends not only on the acoustic features and rate of vocalizations, but also on their timing relative to the vocalizations of others. In this study, we examined whether ultrasonic vocalizations (USVs) produced by adult female Mongolian gerbils during freely moving social interactions are temporally related to the vocal timing of a partner. Using USVCAM, we assigned each USV to an individual caller and evaluated whether brief USVs exhibit turn-taking-like temporal organization by analyzing response latency after partner calls, call overlap rate, and circular-shift surrogate data. The offset-to-onset gaps of alternating vocalizations were concentrated within a short time range, with a median of 130 ms. In addition, the call overlap rate was significantly lower in the observed data than in the circularly shifted surrogate data, indicating that USVs occurred with short latencies after partner calls while being less likely to overlap with ongoing partner calls. Furthermore, USV counts were reduced in unoperated Ctrl animals that interacted with devocalized Mute animals. These findings suggest that gerbil USVs may be coordinated in relation to partner vocal timing and vocal input. This study provides a basis for understanding rodent USVs not merely as individual vocal outputs, but as temporally organized dyadic social signals.

## 1. Introduction

Vocal communication is shaped not only by what acoustic signals are produced and how often they occur, but also by when they are produced relative to the vocalizations of others. In human conversation, speakers alternate their utterances on a short time scale and coordinate speech timing to minimize both silent gaps and overlapping speech.^1,2^ Such temporal coordination is not unique to humans; it has also been investigated in non-human animals as vocal turn-taking or turn-taking-like vocal coordination. Understanding vocal timing coordination in animals is essential for determining how vocal signals are temporally organized within social interactions, rather than being treated merely as isolated vocal outputs.

When evaluating turn-taking-like vocal coordination in animals, response latency after partner calls, call alternation, and avoidance of call overlap are commonly treated as key temporal features.^3,4^ For example, in common marmosets, the timing of vocal exchanges during phee-call antiphonal calling has been shown to have behavioral significance, and in meerkat group calling, calls are socially elicited while overlap with calls from other individuals is avoided.^5,6^ Vocal timing coordination between individuals has also been reported in avian duets and vocal exchanges.^7,8^ These findings indicate that vocal timing can constitute a form of temporal organization that is adjusted in relation to a partner’s calls, rather than being merely a byproduct of vocalization rate. However, because these phenomena should not be equated with human conversation, turn-taking-like temporal organization needs to be evaluated carefully in relation to the social context, call type, and time scale of each species.

In rodents, vocal signals are also closely associated with social interactions and emotional states. In particular, ultrasonic vocalizations (USVs) in rats and mice have been studied in a wide range of contexts, including social approach, sexual behavior, play, mother–infant interactions, and the communication of emotional states.^9–11^ As a well-characterized example of vocal timing control in rodents, male Alston’s singing mice (*Scotinomys teguina*) have been reported to adjust their vocal timing on a subsecond time scale in response to a partner’s song during social interactions.^12^ However, this example involves vocal exchanges based on specialized, long song sequences, which differ in call type and time scale from the brief USVs commonly analyzed in rats, mice, and other laboratory rodents.

Recent advances in sound-source localization have made it possible to assign brief USVs produced during freely moving social interactions in laboratory rodents to individual callers. In laboratory mice, caller assignment using a microphone array has shown that not only males but also females produce USVs during courtship interactions.^13^ In addition, USVCAM has enabled the localization and caller assignment of USV events, including temporally overlapping calls, and has revealed vocal interactions in female ICR mice during resident–intruder encounters.^14^ However, caller assignment or the description of vocal interactions does not, by itself, demonstrate turn-taking-like temporal organization. In particular, few studies have explicitly examined both short-latency responses to partner calls and avoidance of call overlap in brief USVs produced during close-range freely moving social interactions.

Mongolian gerbils (*Meriones unguiculatus*) are social rodents with a diverse vocal repertoire and produce USVs during adult social interactions.^15,16^ Thus, Mongolian gerbils provide a useful model for examining the temporal structure of brief USVs produced during close-range freely moving social interactions. In particular, determining whether gerbil USVs occur as short-latency responses to partner calls and are less likely to overlap with partner calls is important for understanding rodent USVs not merely in terms of individual call output or acoustic features, but as temporally organized dyadic social signals.

In this study, we examined whether USVs produced by adult female Mongolian gerbils during freely moving social interactions are temporally related to the vocal timing of a partner. Using USVCAM, we assigned each USV to an individual caller and analyzed response latency after partner calls and call overlap to evaluate whether brief USVs exhibit turn-taking-like temporal organization. We further examined whether vocal input from a partner is associated with USV production in intact animals by using interactions with devocalized animals. Together, these analyses allowed us to assess whether gerbil USVs can be understood not merely as individual vocal outputs, but as temporally organized dyadic social signals.

## 2. Methods

### 2.1. Animals

All procedures were conducted in accordance with the animal experiment guidelines of Doshisha University. Three-month-old female Mongolian gerbils (*Meriones unguiculatus*) were used in this study. The animals were derived from a laboratory colony originally obtained from Shimizu Laboratory Supplies Co., Ltd. (Kyoto, Japan) and bred and maintained in our laboratory. All animals were sexually naive females and were group-housed with littermates. Animals were housed in standard plastic cages containing paper bedding (Clean Chip SP, Shimizu Laboratory Supplies Co., Ltd., Kyoto, Japan). The animal room was maintained under a 12-h light/dark cycle, with lights on from 08:00 to 20:00. Room temperature was maintained at 22 ± 1°C, relative humidity was maintained at 50–60%, and food and water were available ad libitum.

In Experiment 1, 20 animals were used to form 10 pairs of unfamiliar females. Each animal was used only once. Pairs were composed of animals from different home cages, and pairings between littermates were avoided. None of the animals had previous direct contact experience with non-littermates. Pairs were formed randomly, but the within-pair body weight difference was confirmed to be less than 10 g to reduce the likelihood of aggressive behavior.

In Experiment 2, we used 10 Ctrl × Ctrl pairs consisting of two females, eight Mute × Ctrl pairs consisting of a devocalized animal that had undergone left recurrent laryngeal nerve transection and an unoperated control animal, and eight Sham × Ctrl pairs consisting of a sham-operated animal and an unoperated control animal. The 20 animals in the 10 Ctrl × Ctrl pairs were the same animals used in Experiment 1. The Ctrl animals in the Mute × Ctrl and Sham × Ctrl pairs were unoperated and unfamiliar with the operated animals. These animals were also 3-month-old sexually naive females and were maintained under the same housing conditions as those used in Experiment 1. The pairing criteria for Experiment 2 were the same as those for Experiment 1.

### 2.2. Apparatus and recording conditions

Behavior and USVs were recorded in a plastic arena measuring 30 × 30 × 60 cm (width × depth × height). Fresh paper bedding was placed on the floor of the arena before each trial. Between trials, the arena was cleaned with 70% ethanol. Experiments were conducted in a sound-attenuated room under room lighting, and illumination was kept consistent across trials.

Audio and video recordings were obtained using USVCAM (Katou Acoustics Consultant Office, Kanagawa, Japan). The USVCAM system consisted of an eight-channel MEMS microphone array arranged in a circular configuration, an eight-channel microphone amplifier, an eight-channel high-speed A/D converter, and an imaging device (RealSense D435, Intel Corporation, Santa Clara, CA, USA). Audio was recorded at a sampling rate of 384 kHz with 16-bit resolution. Video was recorded at 30 frames per second. The USVCAM system was positioned 60 cm above the arena floor. Sound-source localization and caller assignment using USVCAM were performed based on the principles described by Matsumoto et al.^14^

### 2.3. Experiment 1: recording of social interactions between female pairs

On the day before the experiment, all animals were fitted with bibs to enable individual identification and to improve the accuracy of pose estimation using DeepLabCut.^17,18^ The preparation and attachment of the bibs were based on Endo et al.^19^ After bib attachment, each animal was singly housed overnight until the start of the experiment; this period served as habituation to the bib. On the day of the experiment, each animal was habituated to the experimental room for 30 min before recording.

After habituation, two unfamiliar female Mongolian gerbils were placed simultaneously in a novel arena. Behavior and USVs were recorded for 10 min immediately after introduction. During recording, no manipulation was performed that would interfere with freely moving social interactions between the animals. No severe aggression meeting the criterion for terminating the experiment was observed, and no pairs were excluded.

### 2.4. Experiment 2: recording of social interactions with devocalized animals

Animals in the Mute group underwent left recurrent laryngeal nerve transection to restrict vocal production. For surgery, anesthesia was induced with isoflurane, followed by intraperitoneal administration of a mixture of medetomidine, midazolam, and butorphanol. The anesthetic mixture was prepared to provide medetomidine at 0.3 mg/kg, midazolam at 4.0 mg/kg, and butorphanol at 5.0 mg/kg. Hair over the surgical site was shaved, and the area was disinfected with 70% ethanol. After a neck incision was made, the left recurrent laryngeal nerve was exposed around the trachea, and a ~1-mm segment of the nerve was removed. The incision was then sutured. In the Sham group, to control for the effects of the surgical procedure itself, the left recurrent laryngeal nerve was exposed and identified but not transected, before the incision was sutured. In both the Mute and Sham groups, lidocaine was applied during the neck incision and after suturing. After surgery, atipamezole (3.0 mg/kg) was administered intraperitoneally to facilitate recovery from anesthesia. Animals were kept warm for approximately 30 min after surgery and were then singly housed. No animals were excluded because of poor postoperative condition.

On postoperative day 2, each animal that had undergone left recurrent laryngeal nerve transection was placed with an unfamiliar 3-month-old sexually naive female that was not used in the main experiment, and vocalizations were recorded for 10 min using USVCAM. Two animals in which residual vocalizations were detected during this confirmation trial were excluded from the analysis, and the eight animals with a USV count of zero were used as the Mute group in the main experiment. The same confirmation trial was also performed for Sham animals to confirm that vocal production was preserved after surgery. These confirmation trials and the partner animals used in the confirmation trials were not included in the main analysis.

On postoperative day 3, the social interaction test was conducted. The experimental procedure was the same as in Experiment 1. Before recording, each animal was habituated to the experimental room for 30 min, after which two unfamiliar animals were placed simultaneously in a novel arena. Behavior and USVs were recorded for 10 min immediately after introduction. During recording, no manipulation was performed that would interfere with freely moving social interactions between the animals.

### 2.5. USV detection and caller assignment

USVs were detected using USVSEG.^20^ For detection with USVSEG, the target frequency range was set to 20–60 kHz, the minimum duration to 10 ms, the maximum duration to 1.0 s, and the event-splitting threshold to 30 ms. The FFT size was set to 512, and the time step was set to 0.5 ms. After automated detection with USVSEG, all USV events were visually inspected on spectrograms, and non-vocal sounds that were likely caused by animal movements, such as bedding noise, were excluded as noise. In addition, based on the vocal classification described by Kobayasi and Riquimaroux,^15^ we confirmed that the USVs included in the present analysis corresponded to bUFMs. Noise exclusion was performed by one analyst before caller assignment.

Using the recorded videos, body parts of each animal were tracked with DeepLabCut version 2.3.8.^17,18^ Seven body points were tracked: the nose, left and right ears, neck, tail base, and two points that divided the body axis between the neck and tail base into three equal segments. The DeepLabCut model was generated by manually labeling a total of 100 frames, consisting of 50 frames extracted from each of two videos. Of the labeled frames, 95% were used for training and 5% for model evaluation, and the model was trained using ResNet-50. Nose coordinates were used for subsequent caller assignment analyses. In the tracking results, points with likelihood values below 0.8 were excluded. Missing frames were linearly interpolated, and obvious tracking errors were manually corrected without reference to vocalization information.

For synchronization of audio and video data, sound-source localization, and caller assignment, we used the Python scripts available in the GitHub repository associated with Matsumoto et al.^14^ Caller assignment was performed using sound-source localization information from the USVCAM system and the nose coordinates of each animal obtained with DeepLabCut. An example of caller assignment is shown in Fig. 1. USV events were excluded from analyses based on caller identity if sound-source localization failed, if the estimated sound-source location was outside the arena or in an implausible position, or if the caller could not be uniquely assigned by the USVCAM system because the animals overlapped in the overhead camera image, making it difficult to identify nose coordinates or animal positions.

**Fig. 1.**
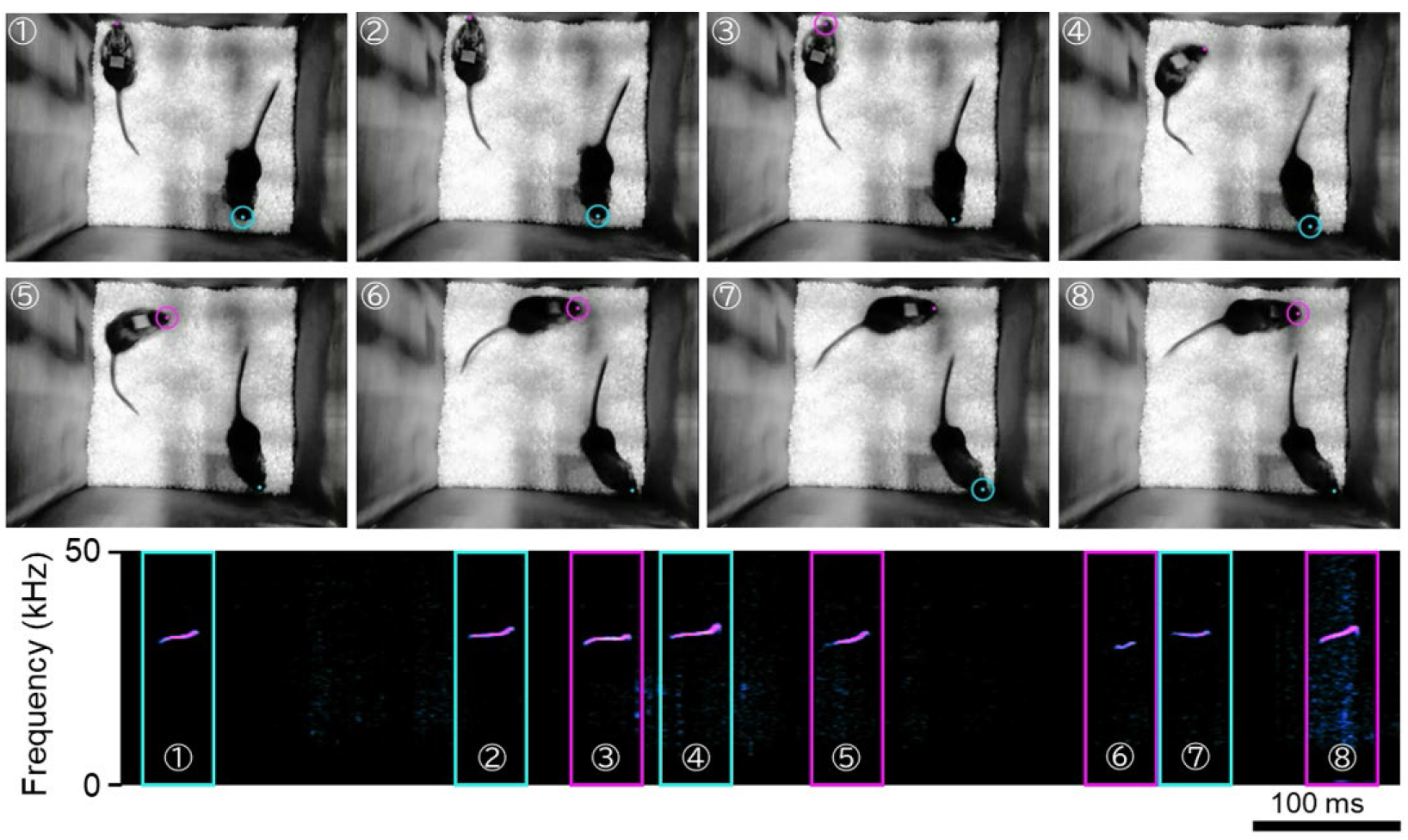
Caller assignment during freely moving social interactions using USVCAM. Two female Mongolian gerbils were recorded with an overhead camera during freely moving social interactions, and the sound-source location of each USV was estimated using USVCAM. The upper panels show video frames at the time points when USVs were detected. Circles indicate sound-source locations estimated by USVCAM, and colors indicate the caller identity. The lower panels show spectrograms of USVs recorded during the same time period. Colored boxes indicate the caller to which each USV was assigned and correspond to the colors of the circles in the upper panels. Circled numbers indicate the correspondence between the video frames in the upper panels and each USV event in the spectrograms in the lower panels. The scale bar indicates 100 ms

### 2.6. Vocal timing analysis

In Experiment 1, because vocalizations were concentrated immediately after recording onset within the 10-min recording period, the first 2 min after recording onset were used for call alternation analysis. For each USV event, the time interval from the offset of that event to the onset of the next USV event was calculated as the offset-to-onset gap. When a subsequent USV event was detected, that event was then used as a new reference call, and the same procedure was repeated. Call transitions between the two animals were analyzed in both directions so that the analysis did not depend on the order of individual IDs. Specifically, alternating vocalizations were defined as cases in which a partner’s call began within 1 s after the offset of the reference call. In contrast, within-animal sequential vocalizations were defined as cases in which the same animal produced the next call within 1 s after the offset of the reference call. Calls separated by intervals of 1 s or longer were not treated as sequential call transitions. Calls were defined as overlapping when the call durations of the two animals overlapped at least partially. Overlapping calls were not included in alternating vocalizations or within-animal sequential vocalizations, but were analyzed separately as a proportion of all USV events.

To examine whether the temporal structure of the observed call transitions deviated from the distribution expected by chance, we performed a circular-shift analysis. For each pair, the call sequence of one animal was fixed, and the call sequence of the other animal was randomly circularly shifted within the 2-min analysis window. This procedure was performed for call transitions in both directions. The shift amount was randomly selected from 0 to 120 s, and 1,000 circular shifts were performed for each pair. The duration of each USV event was preserved after shifting. When a shifted USV event extended beyond the end of the analysis window, the 2-min analysis window was treated as a circular time axis, and the boundary-crossing USV event was split between the end and beginning of the window. For each circularly shifted dataset, offset-to-onset gaps were calculated using the same criteria as those used for the observed data.

In Experiment 2, to examine whether the number of USVs produced by unoperated Ctrl animals varied depending on partner condition, the total number of USVs produced by each animal during the entire 10-min recording period was calculated.

### 2.7. Statistical analysis

Statistical analyses were performed using GraphPad Prism version 10 (GraphPad Software LLC, Boston, MA, USA). In Experiment 1, statistical analyses were performed using pairs as the unit of analysis. The call overlap rate was calculated as the proportion of overlapping calls relative to the total number of USV events in each pair. The call overlap rate in the observed data was compared with that in the surrogate data obtained by circular-shift analysis. The Wilcoxon matched-pairs signed-rank test was used to compare the observed and circularly shifted data.

In Experiment 2, we examined whether the number of USVs produced by unoperated Ctrl animals varied depending on pair type. In Ctrl × Ctrl pairs, the mean number of USVs produced by the two animals was used as the representative value for each pair. In Mute × Ctrl and Sham × Ctrl pairs, the number of USVs produced by the unoperated Ctrl animal in each pair was used for analysis. The three groups were compared using the Kruskal–Wallis test, followed by Dunn’s multiple comparisons test for post hoc comparisons. The significance threshold was set at p < 0.05. Each point in the figures for Experiments 1 and 2 represents one pair.

## 3. Results

### 3.1. Experiment 1: temporal structure of vocal timing

#### 3.1.1. Gap distributions of alternating vocalizations and within-animal sequential vocalizations

Two unfamiliar female Mongolian gerbils were allowed to interact freely in a novel arena, and USVs were recorded and assigned to individual callers using USVCAM. Based on the vocal classification described by Kobayasi and Riquimaroux,^15^ the USVs recorded under the present conditions corresponded to bUFMs. Under these conditions, we observed cases in which one animal produced a USV shortly after a call from its partner. When the timing of the next calls was aligned to the offset of the reference call, next calls were frequently distributed immediately after the offset of the reference call (Fig. 2a). Similarly, when the timing of the next calls was aligned to the onset of the reference call, next calls did not simply overlap randomly immediately after the onset of the reference call, but were more frequently observed after the offset of the reference call (Fig. 2b). Focusing on the temporal overlap between partner calls and subsequent calls, next calls were more frequently distributed after the offset of partner calls than during partner calls. Thus, under the present conditions, USVs showed a temporal structure in which calls were more likely to occur after the offset of partner calls rather than overlapping with ongoing partner calls.

**Fig. 2.**
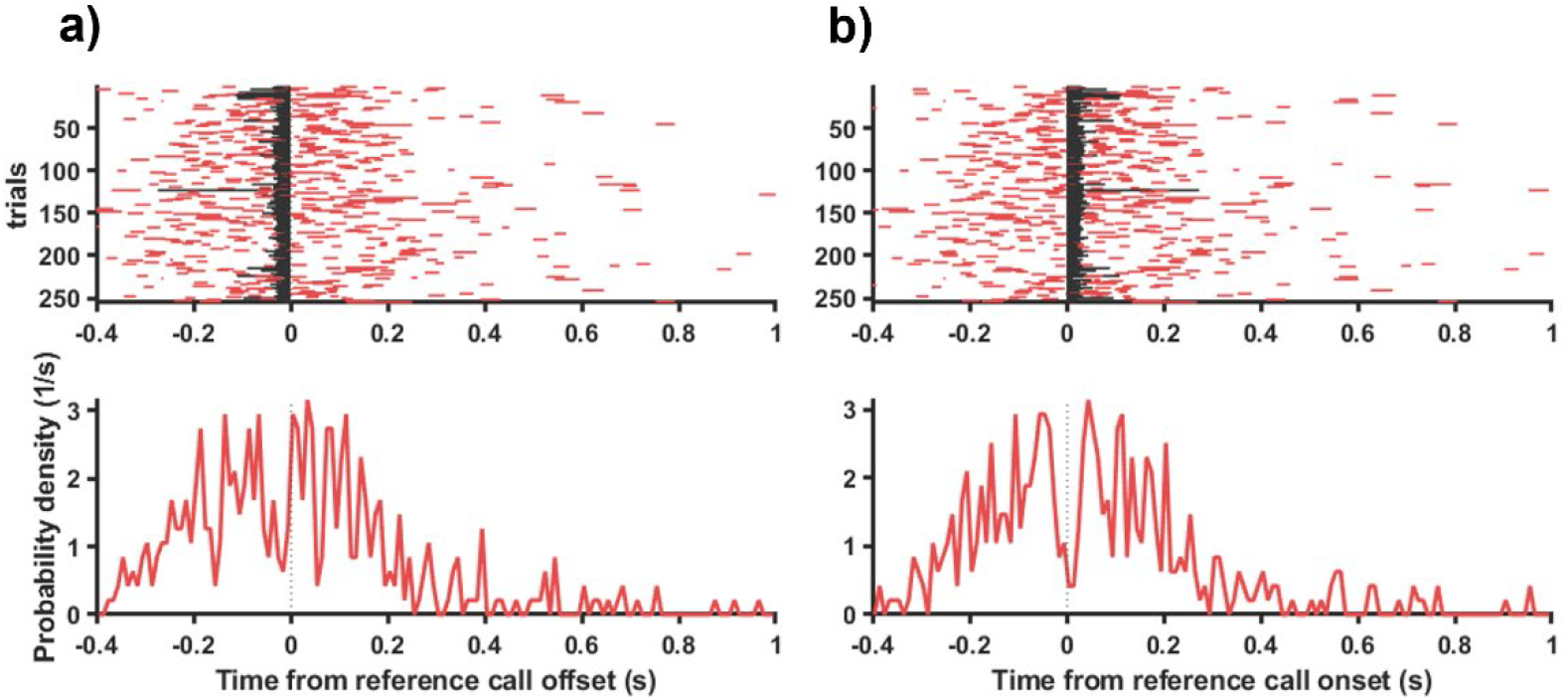
Distribution of vocal timing relative to partner calls. For USVs recorded during freely moving social interactions between two unfamiliar female Mongolian gerbils, calls produced by one animal were treated as reference calls, and the timing of calls produced by the partner was aligned relative to them. The upper panels show raster plots, and the lower panels show probability density distributions of vocal timing. Red lines indicate partner call events, and black lines indicate the duration of the reference call. (a) Distribution aligned with the offset of the reference call set to 0 s. (b) Distribution aligned with the onset of the reference call set to 0 s

Next, we quantitatively evaluated the temporal structure of alternating vocalizations by calculating the time interval from the offset of one animal’s call to the onset of the next call by the partner as the offset-to-onset gap. The offset-to-onset gaps of alternating vocalizations were concentrated within a short time range, with a median of approximately 130 ms (IQR = 70–227 ms; Fig. 3a). This result indicates that, during social interactions in Mongolian gerbils, call transitions occurred in which a partner produced a call shortly after one animal’s call. To examine whether the short gaps observed in alternating vocalizations reflected between-animal responsiveness or simply the tendency of each animal’s call sequence to contain short-interval calls, we also calculated offset-to-onset gaps for within-animal sequential vocalizations. In within-animal sequential vocalizations, gaps were also concentrated within a short time range (Fig. 3b). This finding indicates that call transitions with short gaps were not limited to alternating vocalizations, but were also present within each animal’s own call sequence. Therefore, it was necessary to determine whether the short gaps observed in alternating vocalizations were attributable merely to short-interval structure within each animal’s own call sequence or reflected temporal correspondence in vocal timing between the two animals. To address this issue, we generated surrogate data by fixing the call sequence of one animal and randomly circularly shifting the call sequence of the other animal, and compared these data with the observed data.

**Fig. 3.**
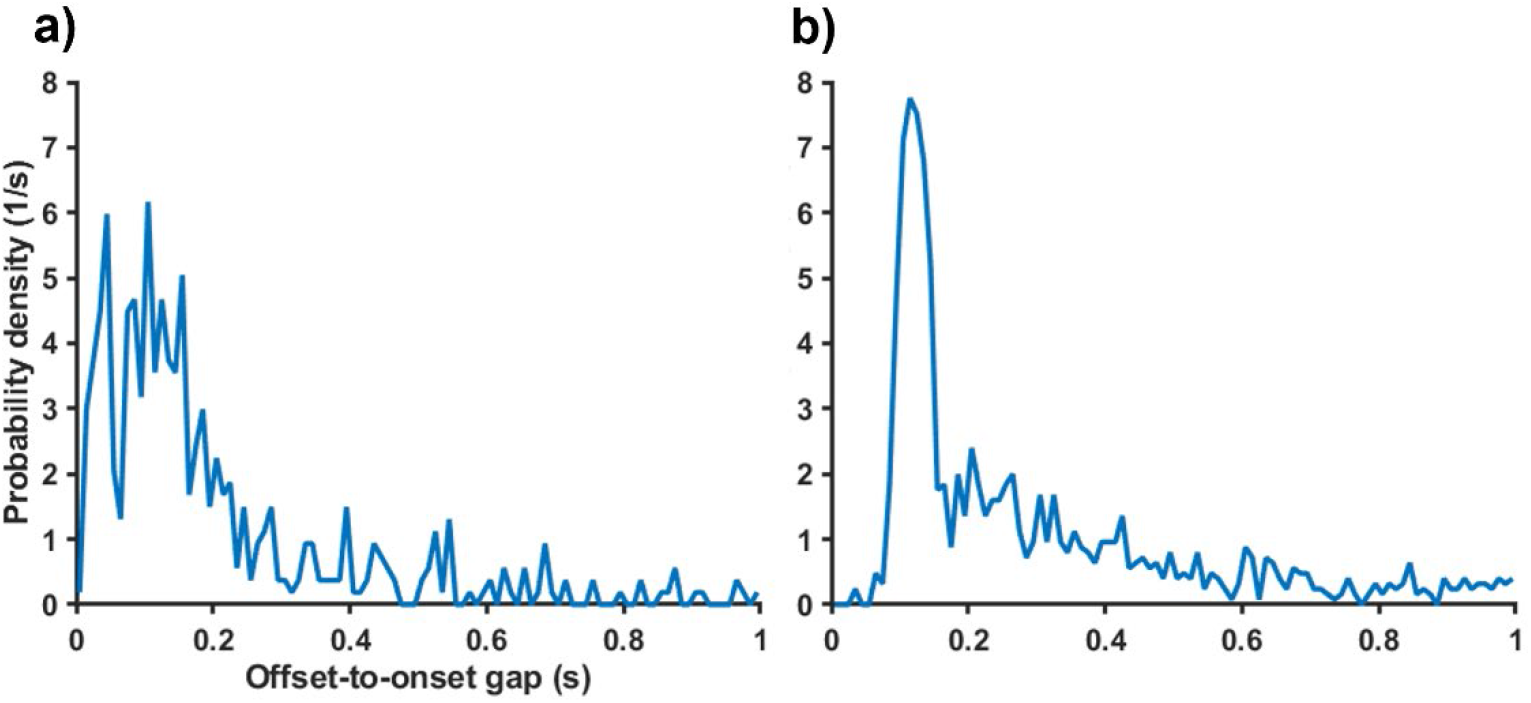
Offset-to-onset gap distributions of alternating vocalizations and within-animal sequential vocalizations. For USVs recorded during freely moving social interactions between female pairs, the time interval from call offset to the onset of the next call was calculated as the offset-to-onset gap, and the probability density distributions are shown. (a) Gap distribution of alternating vocalizations, defined as cases in which a partner call began within 1 s after the offset of the reference call. (b) Gap distribution of within-animal sequential vocalizations, defined as cases in which the same animal produced the next call within 1 s after the offset of the reference call

#### 3.1.2. Circular-shift analysis of non-random vocal timing

In the observed data, the gap distribution of alternating vocalizations showed a concentration of short gaps, with a median of approximately 130 ms. In contrast, this concentration was weaker in the circularly shifted surrogate data, which showed a broader distribution (Fig. 4a). These results suggest that the concentration of short gaps in alternating vocalizations was difficult to explain solely by the tendency of each animal to produce sequential calls at short intervals, and may reflect temporal correspondence in vocal timing between the two animals.

**Fig. 4.**
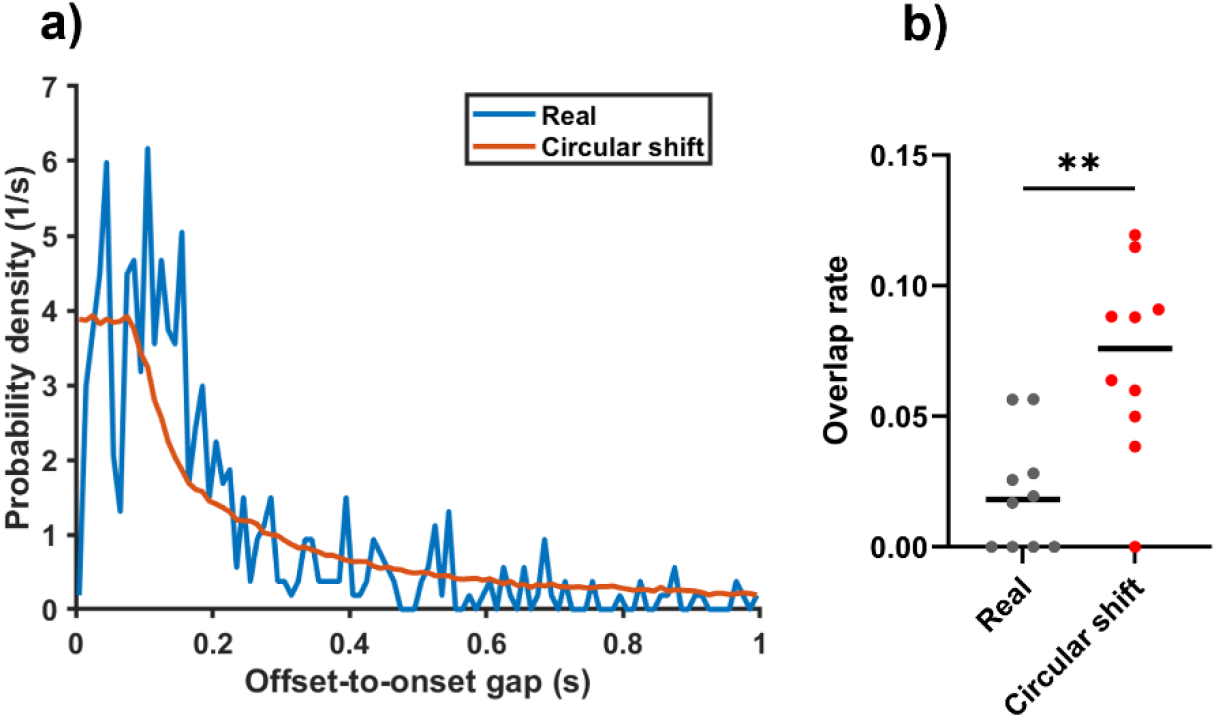
Comparison of alternating-vocalization gap distributions and call overlap rates using circular-shift analysis. For USVs recorded during freely moving social interactions between female pairs, surrogate data were generated by fixing the call sequence of one animal and randomly circularly shifting the call sequence of the other animal, and these were compared with the observed data. (a) Offset-to-onset gap distributions of alternating vocalizations in the observed data and the circularly shifted surrogate data. The blue line indicates the observed data, and the orange line indicates the circularly shifted surrogate data. (b) Call overlap rates in the observed data and the circularly shifted surrogate data. Each point represents one pair. For the circularly shifted surrogate data, the mean call overlap rate obtained from 1,000 circular shifts for each pair is shown as the representative value for that pair. Horizontal lines indicate medians. \*\**p* < 0.01

We further compared the call overlap rate between the observed data and the circularly shifted surrogate data. For the circularly shifted surrogate data, the mean call overlap rate obtained from 1,000 circular shifts for each pair was used as the representative value for that pair. The call overlap rate was significantly lower in the observed data than in the circularly shifted surrogate data (Wilcoxon matched-pairs signed-rank test, *N* = 10 pairs, *W* = 45.0, *p* = 0.00390; Fig. 4b). This result indicates that gerbil USVs were likely to occur with short gaps immediately after partner calls, while being less likely to overlap with ongoing partner calls. Together, these findings indicate that USVs produced during social interactions between female Mongolian gerbils showed a temporal structure characterized by short-latency occurrence after partner calls and reduced call overlap.

### 3.2. Experiment 2: partner vocal input and USV production in unoperated Ctrl animals

In Experiment 1, short-latency alternating vocalizations and reduced call overlap were observed in pairs of intact females in relation to partner vocal timing. Therefore, in Experiment 2, to examine whether such vocalizations were related to vocal input from the partner, we used Mute animals whose vocal production had been restricted by left recurrent laryngeal nerve transection and analyzed vocalizations during social interactions with unoperated Ctrl animals. In postoperative confirmation trials, no USVs were detected in the animals used as the Mute group. In contrast, vocal production was preserved in Sham-operated animals. These results confirmed that recurrent laryngeal nerve transection restricted vocal production, whereas the sham surgical procedure alone did not substantially impair the ability to produce vocalizations.

We next examined whether USV counts in unoperated Ctrl animals varied depending on the vocal ability of the partner animal. In Ctrl × Ctrl pairs, the mean number of USVs produced by the two animals was used as the representative value for each pair. In Mute × Ctrl and Sham × Ctrl pairs, the number of USVs produced by the unoperated Ctrl animal in each pair was used. Comparison among the three groups showed that USV counts differed depending on partner condition (Kruskal– Wallis test, *H* = 15.6, *df* = 2, *N* = 26, *p* < 0.001; Fig. 5). Post hoc comparisons showed that USV counts in unoperated Ctrl animals paired with Mute animals were significantly lower than those in Ctrl × Ctrl pairs (Dunn’s multiple comparisons test, *p* = 0.0178). USV counts in unoperated Ctrl animals paired with Mute animals were also significantly lower than those in unoperated Ctrl animals paired with Sham animals (*p* < 0.001). In contrast, no significant difference was observed between Ctrl animals in Ctrl × Ctrl pairs and those in Sham × Ctrl pairs (*p* = 0.563). These results indicate that the reduction in USV counts in unoperated Ctrl animals was associated with the absence of vocal input from the partner animal, rather than with the presence of an operated partner per se. Together, these findings suggest that the absence of vocal input from the partner animal is associated with reduced USV production in unoperated Ctrl animals during social interactions.

**Fig. 5.**
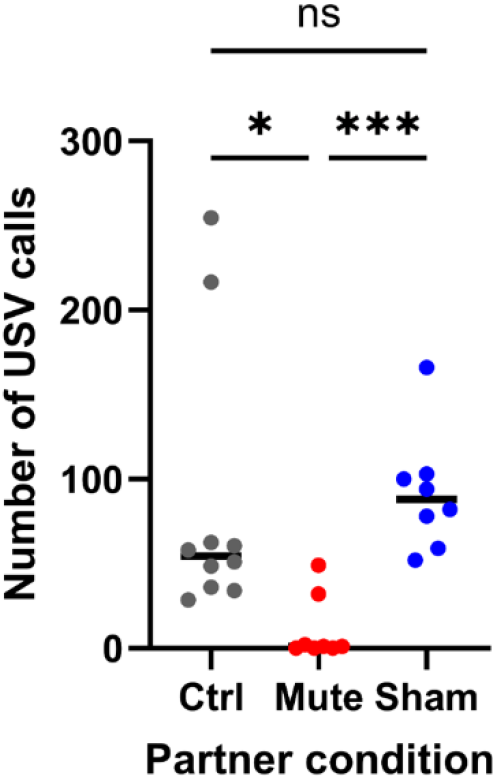
USV counts in unoperated Ctrl animals paired with Ctrl, Mute, or Sham animals. The number of USVs recorded from unoperated Ctrl animals during 10 min of freely moving social interactions is shown for Ctrl × Ctrl, Mute × Ctrl, and Sham × Ctrl pairs. In Ctrl × Ctrl pairs, the mean number of USVs produced by the two animals was used as the representative value for each pair. In Mute × Ctrl and Sham × Ctrl pairs, the number of USVs produced by the unoperated Ctrl animal in each pair is shown. Each point represents one pair, and horizontal lines indicate medians. \**p* < 0.05, \*\*\**p* < 0.001.

## 4. Discussion

This study showed that USVs produced by adult female Mongolian gerbils were temporally related to partner calls during freely moving social interactions. Caller assignment using USVCAM revealed that calls by the partner were frequently observed with short latencies after the offset of one animal’s call (Figs. 2 and 3), and that call overlap was lower in the observed data than in the circularly shifted surrogate data (Fig. 4b). In addition, USV counts were reduced in unoperated Ctrl animals paired with Mute animals compared with those in the Ctrl × Ctrl and Sham × Ctrl conditions (Fig. 5). These findings suggest that gerbil USVs are not produced merely as independent vocal outputs reflecting each animal’s own vocal tendency, but may be coordinated in relation to partner vocal timing and vocal input. Thus, this study provides evidence for turn-taking-like temporal coordination of USVs during close-range social interactions in Mongolian gerbils.

The median offset-to-onset gap of alternating vocalizations observed in this study was 130 ms, with an interquartile range of 70–227 ms (Fig. 3a). In addition, the gap distribution of alternating vocalizations in the observed data showed a clear peak within a short time range after partner call offset, whereas this concentration was weaker in the circularly shifted surrogate data (Fig. 4a). These results indicate that gerbil USV exchanges are organized on a short time scale of several hundred milliseconds. In marmoset antiphonal calling, vocal responses have been reported to occur on a time scale of several seconds.^5^ In contrast, the gap between conversational turns in humans is generally considered to be approximately 200 ms.^1,2^ Thus, the short-latency USV exchanges observed in the present study may operate on a time scale distinct from long-range antiphonal calling with marmoset phee calls. However, because marmoset phee calls and gerbil USVs differ substantially in call duration, social context, and inter-individual distance, differences in gap duration should not be interpreted directly as differences in the function or control mechanisms of vocal turn-taking. Rather, the short gaps observed in this study should be interpreted as a finding that characterizes the time scale of brief USV exchanges during close-range freely moving social interactions.

However, call transitions with short gaps alone cannot rule out the alternative interpretation that the two animals were in a high-vocalization state induced by the novel environment or social contact and were simply producing calls sequentially at short intervals. Indeed, short gaps were also present in within-animal sequential vocalizations in the present study (Fig. 3b). What is important in this regard is that the call overlap rate in the observed data was lower than that in the circularly shifted surrogate data (Fig. 4b). Because circular-shift analysis disrupts the temporal relationship between the two animals while preserving each animal’s call sequence, the concentration of short gaps and reduced call overlap seen in the observed data are difficult to explain solely by call density or a tendency to produce short-interval call sequences. In meerkat group calling, socially elicited calling combined with avoidance of overlap with calls from other individuals has also been reported as an important feature of turn-taking.^6^ Therefore, the USV exchanges observed in the present study can be interpreted as a turn-taking-like dyadic vocal structure characterized by both a short-gap time scale and reduced call overlap.

Such vocal timing coordination highlights that gerbil vocal communication should be considered not only in terms of vocal repertoire or call rate, but also in terms of dyadic temporal structure within a single call type. Adult Mongolian gerbils have a diverse vocal repertoire, and Kobayasi and Riquimaroux^15^ classified their vocalizations into multiple syllable types. The USVs analyzed in the present study corresponded to bUFMs in this classification. Thus, these findings highlight the importance of considering how calls of the same type are temporally placed relative to partner calls during social interactions.

Vocal input from the partner may be a factor that supports USV production during social interactions. Unoperated Ctrl animals paired with Mute animals showed lower USV counts than those in the Ctrl × Ctrl and Sham × Ctrl conditions (Fig. 5). Because vocal production was preserved in Sham animals, this reduction may be associated with the absence of vocal input from the partner, rather than with the presence of an operated partner per se. Studies using devocalization in rodents have shown that intact animals can continue to vocalize even when their partner cannot produce vocalizations, but the effects of devocalization appear to depend on the social context. In the sexual behavior context in rats, intact animals interacting with a devocalized partner have been reported to produce USVs, and devocalization does not substantially alter male–female copulatory behavior.^21^ In contrast, in the play context in rats, play is reduced in devocalized pairs, suggesting that 50-kHz USVs may contribute to the facilitation or maintenance of playful interactions.^22^ Taken together, these findings suggest that the extent to which vocal input from a partner contributes to the production of an animal’s own vocalizations may depend not only on species differences, but also on the social context and call type. The present findings suggest that, during close-range interactions between adult female Mongolian gerbils, USV input from the partner may support USV production in unoperated Ctrl animals. This provides an important basis for examining the social feedback function of rodent USVs.

However, the results shown in Fig. 5 do not demonstrate that the absence of vocal input from the partner directly caused the reduction in USV counts in unoperated Ctrl animals. Although the inclusion of the Sham group controlled, to some extent, for the effects of neck surgery and postoperative experience, recurrent laryngeal nerve transection is not a method that selectively manipulates only vocal input from the partner. Therefore, caution is needed when attributing the reduction in USV counts in unoperated Ctrl animals solely to the absence of vocal input from the partner. Accordingly, the present results should be interpreted as showing that the vocal ability of the partner animal is associated with USV production in unoperated Ctrl animals during close-range interactions in Mongolian gerbils. Future playback experiments that manipulate the presence or timing of partner calls will allow a more direct test of the causal role of partner vocal input in maintaining an animal’s own vocalizations.

The turn-taking-like temporal organization shown in the present study was based on USV exchanges immediately after unfamiliar adult females began interacting in a novel environment. In addition, the call alternation analysis focused on the first 2 min after recording onset, during which vocalizations were concentrated. Therefore, determining how this temporal organization changes across different social contexts or over the time course of social interactions will be an important direction for future studies.

In conclusion, this study showed that USVs corresponding to bUFMs in adult female Mongolian gerbils were temporally related to partner calls during freely moving social interactions. These USVs occurred with short latencies after partner calls and were less likely to overlap with ongoing partner calls. In addition, USV counts decreased in unoperated Ctrl animals when the partner animal lacked vocal ability, suggesting that USV production in Mongolian gerbils may be supported by vocal input from the partner. These findings indicate that rodent USVs should be understood not only in terms of individual call output or acoustic features, but also as temporally organized dyadic social signals. This study provides a basis for understanding vocal communication in Mongolian gerbils during close-range social interactions from the perspective of turn-taking-like temporal coordination.

## Acknowledgements

We thank Prof. Shizuko Hiryu, Dr. Ryosuke O. Tachibana, and Dr. Masahiro Kato for their valuable suggestions.

## References

1. Stivers, T., Enfield, N.J., Brown, P., Englert, C., Hayashi, M., Heinemann, T., Hoymann, G., Rossano, F., de Ruiter, J.P., Yoon, K.-E., et al. (2009). Universals and cultural variation in turn-taking in conversation. Proc. Natl. Acad. Sci. U. S. A. 106, 10587–10592. 10.1073/pnas.0903616106.

2. Levinson, S.C., and Torreira, F. (2015). Timing in turn-taking and its implications for processing models of language. Front. Psychol. 6, 731. 10.3389/fpsyg.2015.00731.

3. Henry, L., Craig, A.J.F.K., Lemasson, A., and Hausberger, M. (2015). Social coordination in animal vocal interactions. Is there any evidence of turn-taking? The starling as an animal model. Front. Psychol. 6, 1416. 10.3389/fpsyg.2015.01416.

4. Pika, S., Wilkinson, R., Kendrick, K.H., and Vernes, S.C. (2018). Taking turns: bridging the gap between human and animal communication. Proc. Biol. Sci. 285, 20180598. 10.1098/rspb.2018.0598.

5. Miller, C.T., Beck, K., Meade, B., and Wang, X. (2009). Antiphonal call timing in marmosets is behaviorally significant: interactive playback experiments. J. Comp. Physiol. A Neuroethol. Sens. Neural Behav. Physiol. 195, 783–789. 10.1007/s00359-009-0456-1.

6. Demartsev, V., Strandburg-Peshkin, A., Ruffner, M., and Manser, M. (2018). Vocal turn-taking in meerkat group calling sessions. Curr. Biol. 28, 3661–3666.e3. 10.1016/j.cub.2018.09.065.

7. Fortune, E.S., Rodríguez, C., Li, D., Ball, G.F., and Coleman, M.J. (2011). Neural mechanisms for the coordination of duet singing in wrens. Science 334, 666–670. 10.1126/science.1209867.

8. Benichov, J.I., and Vallentin, D. (2020). Inhibition within a premotor circuit controls the timing of vocal turn-taking in zebra finches. Nat. Commun. 11, 221. 10.1038/s41467-019-13938-0.

9. Brudzynski, S.M. (2013). Ethotransmission: communication of emotional states through ultrasonic vocalization in rats. Curr. Opin. Neurobiol. 23, 310–317. 10.1016/j.conb.2013.01.014.

10. Wöhr, M., and Schwarting, R.K.W. (2013). Affective communication in rodents: ultrasonic vocalizations as a tool for research on emotion and motivation. Cell Tissue Res. 354, 81–97. 10.1007/s00441-013-1607-9.

11. Portfors, C.V., and Perkel, D.J. (2014). The role of ultrasonic vocalizations in mouse communication. Curr. Opin. Neurobiol. 28, 115–120. 10.1016/j.conb.2014.07.002.

12. Okobi, D.E., Jr, Banerjee, A., Matheson, A.M.M., Phelps, S.M., and Long, M.A. (2019). Motor cortical control of vocal interaction in neotropical singing mice. Science 363, 983–988. 10.1126/science.aau9480.

13. Neunuebel, J.P., Taylor, A.L., Arthur, B.J., and Egnor, S.E.R. (2015). Female mice ultrasonically interact with males during courtship displays. Elife 4, e06203. 10.7554/eLife.06203.

14. Matsumoto, J., Kanno, K., Kato, M., Nishimaru, H., Setogawa, T., Chinzorig, C., Shibata, T., and Nishijo, H. (2022). Acoustic camera system for measuring ultrasound communication in mice. iScience 25, 104812. 10.1016/j.isci.2022.104812.

15. Kobayasi, K.I., and Riquimaroux, H. (2012). Classification of vocalizations in the Mongolian gerbil, Meriones unguiculatus. J. Acoust. Soc. Am. 131, 1622–1631. 10.1121/1.3672693.

16. Furuyama, T., Shigeyama, T., Ono, M., Yamaki, S., Kobayasi, K.I., Kato, N., and Yamamoto, R. (2022). Vocalization during agonistic encounter in Mongolian gerbils: Impact of sexual experience. PLoS One 17, e0272402. 10.1371/journal.pone.0272402.

17. Mathis, A., Mamidanna, P., Cury, K.M., Abe, T., Murthy, V.N., Mathis, M.W., and Bethge, M. (2018). DeepLabCut: markerless pose estimation of user-defined body parts with deep learning. Nat. Neurosci. 21, 1281–1289. 10.1038/s41593-018-0209-y.

18. Nath, T., Mathis, A., Chen, A.C., Patel, A., Bethge, M., and Mathis, M.W. (2019). Using DeepLabCut for 3D markerless pose estimation across species and behaviors. Nat. Protoc. 14, 2152‒2176. 10.1038/s41596-019-0176-0.

19. Endo, N., Makinodan, M., Mannari-Sasagawa, T., Horii-Hayashi, N., Somayama, N., Komori, T., Kishimoto, T., and Nishi, M. (2021). The effects of maternal separation on behaviours under social-housing environments in adult male C57BL/6 mice. Sci. Rep. 11, 527. 10.1038/s41598-020-80206-3.

20. Tachibana, R.O., Kanno, K., Okabe, S., Kobayasi, K.I., and Okanoya, K. (2020). USVSEG: A robust method for segmentation of ultrasonic vocalizations in rodents. PLoS One 15, e0228907. 10.1371/journal.pone.0228907.

21. Ågmo, A., and Snoeren, E.M.S. (2015). Silent or vocalizing rats copulate in a similar manner. PLoS One 10, e0144164. 10.1371/journal.pone.0144164.

22. Kisko, T.M., Euston, D.R., and Pellis, S.M. (2015). Are 50-khz calls used as play signals in the playful interactions of rats? III. The effects of devocalization on play with unfamiliar partners as juveniles and as adults. Behav. Processes 113, 113–121. 10.1016/j.beproc.2015.01.016.

